# Quantitative Genetic Analysis of the Maize Leaf Microbiome

**DOI:** 10.1101/268532

**Authors:** Jason G. Wallace, Karl A. Kremling, Edward S. Buckler

## Abstract

The degree to which an organism can affect its associated microbial communities (“microbiome”) varies by organism and habitat, and in many cases is unknown. We address this question by analyzing the metabolically active bacteria of the maize phyllosphere across 300 diverse maize lines growing in a common environment. We performed comprehensive heritability analysis for 49 community diversity metrics, 380 bacterial clades (individual operational taxonomic units and higher-level groupings), and 9042 predicted metagenomic functions. We find that only a few few bacterial clades (5) and diversity metrics (2) are significantly heritable, while a much larger number of metabolic functions (200) are. Many of these associations appear to be driven by the amount of Methylobacteria present in each sample, and we find significant enrichment for traits relating to short-chain carbon metabolism, secretion, and nitrotoluene degradation. Genome-wide association analysis identifies a small number of associated loci for these heritable traits, including two loci (on maize chromosomes 7 and 10) that affect a large number of traits even after correcting for correlations among traits. This work is among the most comprehensive analyses of the maize phyllosphere to date. Our results indicate that while most of the maize phyllosphere composition is driven by environmental factors and/or stochastic founder events, a subset of bacterial taxa and metabolic functions is nonetheless significantly impacted by host plant genetics. Additional work will be needed to identify the exact nature of these interactions and what effects they may have on the phenotype of host plants.

## Introduction

A major question in microbiome science is the degree to which a host organism can affect its associated microbial communities. In animals, several host-associated microbiomes have shown significant effects of host genetics, especially in the gut (Goodrich et al. 2014, 2016; Xie et al. 2016; Gomez et al. 2017; Demmitt et al. 2017). In some cases specific genes have been implicated in these associations (Bonder et al. 2016; Goodrich et al. 2016; Belheouane et al. 2017; Lynch et al. 2017). In plants, host genotype has been implicated in structuring the microbiomes of temperate and tropical forests (Laforest-Lapointe, Messier, and Kembel 2016; Kim et al. 2012; Kembel et al. 2014; Redford et al. 2010), *Arabidopsis thaliana* (Knief et al. 2010; Horton et al. 2014; Lundberg et al. 2012; Bulgarelli et al. 2012), maize (Peiffer et al. 2013), grapevine (Zarraonaindia et al. 2015), tobacco (Knief et al. 2010), wild mustard (Wagner et al. 2016), and others. These studies have generally found significant effects of plant genotype on the associated microbial communities, though these genetic effects are usually overshadowed by environmental effects (Müller et al. 2016; Finkel et al. 2011; Rastogi et al. 2012).

Whereas soil is arguably the most complex microbial habitat on earth (Daniel 2005) and plant rhizosphere communities are only slightly less complex (Mendes, Garbeva, and Raaijmakers 2013), the plant phyllosphere--the collection of above-ground plant surfaces--is comparatively simple (Bodenhausen, Horton, and Bergelson 2013; Delmotte et al. 2009). The phyllosphere environment is actually very harsh, since plant surfaces have little water and nutrients available and are exposed to environmental stresses such as temperature fluctuations and ultraviolet radiation (Vorholt 2012). Most phyllsopheres communities are dominated by a small number of bacterial clades capable of living in this environment, especially the Alphaproteobacteria, Gammaproteobacteria, Bacteroidetes, and Actinobacteria (Vorholt 2012; Steven E. Lindow and Brandl 2003). These organisms subsist off a variety of carbon compounds derived from their plant host, including carbohydrates, amino acids, organic acids, sugar alcohols, isoprenes, and short-chain carbon compounds (Vorholt 2012; Steven E. Lindow and Brandl 2003). Methanol is a particularly important carbon source for bacteria in the family Methylobacteriaceae, which contains many facultative methylotrophs present in the phyllosphere (Vorholt 2012; Knief et al. 2012).

Investigations into how plant genotype can shapes phyllosphere communities have focused mostly in individual gene mutants. Work in *Arabidopsis* indicates that the phyllosphere community can be influenced by jasmonic acid defense pathway (Kniskern, Traw, and Bergelson 2007), cuticle composition (Bodenhausen et al. 2014), and ethylene signalling (Bodenhausen et al. 2014). Attempts to map natural genetic variation that affects the phyllosphere have been more limited, but have identified the gamma-aminobutyric acid pathway in maize (Balint-Kurti et al. 2010) and both defense and cell wall integrity in *Arabidopsis* (Horton et al. 2014). More generally, phyllosphere communities have been shown to relate to disease resistance (Balint-Kurti et al. 2010; Manching, Balint-Kurti, and Stapleton 2014; Agler et al. 2016), and to respond to drought stress (Methe et al. 2017), ultraviolet radiation (Jacobs and Sundin 2001; Kadivar and Stapleton 2003) and nitrogen fertilization (Manching, Balint-Kurti, and Stapleton 2014).

Although several studies have have identified significant impacts of plant genotype on the phyllosphere community (e.g., (Sapkota et al. 2015; Bálint et al. 2013; Lambais, Lucheta, and Crowley 2014)), most of these have been comparisons of a small number of individuals and/or spanning multiple species (*Arabidopsis* being the chief exception; (Horton et al. 2014; Agler et al. 2016)). To investigate the role of host genetics in phyllosphere composition, we sampled the phyllosphere community across 300 diverse maize lines growing in a common environment. We chose maize both because of its high genetic diversity (Chia et al. 2012) and because of its relevance to global agriculture. Over 1 billion metric tons of maize is harvested each year (37% of global cereal production) (Food and Agriculture Organization of the United Nations. 2017), and it covers an estimated 188 million hectares each year--an area nearly the size of Mexico and representing 13% of global arable land (Food and Agriculture Organization of the United Nations. 2017). Given that many serious disease of maize occur in the phyllosphere (Munkvold and White 2016), understanding the forces that shape phyllosphere communities is a first step toward identifying ways to harness those communities for global agriculture.

## Materials and Methods

### Field setup and tissue sampling

The RNA samples for this experiment were taken from a parallel experiment on maize gene expression (Kremling et al. 2018). In brief, 300 members of the extended Goodman Maize Association Panel (Flint-Garcia et al. 2005) were grown on the Musgrave Research Farm in Aurora, New York, in the summer of 2014. Plants were blocked by maturity (5 blocks) and planted with delays to try to synchronize flowering as much as possible. Tissue samples were collected on two separate dates (August 8, 2014 and August 26, 2014); the first sampling collected from all plots that had flowered up to that point, and the second on all that had flowered after the first but before the second. A small number of check plots were sampled on both dates.

Samples were collected from the midpoint of the second leaf down from the top (**Supplemental Figure 1**). A ˜0.5-1 cm wide strip of leaf tissue from the edge of the leaf to the midrib was cut out with scissors, rolled, and placed into a collection tube. Three plants from each plot were bulked into a single tube, which was then flash-frozen in liquid nitrogen. All samples were collected within a 2-hour period. Leaf strips were collected between 11 AM and 1 PM (“day”), and between 11 PM and 1 AM (“night”), avoiding the previously collected plants and edge plants when possible. This separate day and night collection was intended primarily to capture circadian gene expression for Kremling et al. (2018); for this experiment they serve as technical replicates.

### cDNA library preparation and sequencing

RNA was extracted using TRIzol (Invitrogen, Carlsbad, CA) with Direct-zol columns (Zymo Research, Irvine, CA). RNA concentration was determined with QuantiFluor RNA dye (Promega) and concentrations were normalized to 100 ng/μl. RNA was then converted to cDNA using ProtoScript II reverse transcriptase (New England Biolabs) according to manufacturer’s instructions (1 μl template per reaction), except the enzyme amount was cut in half and the incubation time extended to 60 minutes. The RT primer was 1391R (5’-GACGGGCGGTGWGTRCA-3’) (LANE and J 1991). cDNA was then PCR-amplified in a 30 μl reaction using 2x Hot Start Taq DNA Polymerase master mix (New England Biolabs), 0.5 ul cDNA reaction as template, and the chloroplast-discriminating primers 799F (5’-TCGTCGGCAGCGTCAGATGTGTATAAGAGACAGACCMGGATTAGATACCCKG -3’) and 1115R (5’-GTCTCGTGGGCTCGGAGATGTGTATAAGAGACAGAGGGTTGCGCTCGTTG -3’) (Shade, McManus, and Handelsman 2013), which target the V5/V6 region of the gene. (Note that both primers include linker sequenced on the 5’ end to add Illumina adaptors in the next step.) The PCR program was 95° C for 5 minutes, 15 cycles 95° C (30 sec) - 55° C (30 sec) - 72° C (30 sec), and a final extension at 72° C for 5 minutes. The final PCR reaction was diluted 20-fold in a fresh PCR reaction with Nextera barcoded adapters (Illumina); cycle conditions were identical to above except only 8 cycles were performed. 5 μl of each reaction were pooled and purified using a QIAquick PCR Purification Kit (Qiagen). Libraries were submitted to the Genomics Facility at Cornell University for 2×200 paired-end sequencing on an Illumina HiSeq 2500. Raw sequence data is available through the NCBI Short Read Archive (SRP132042).

### OTU calling

Paired-end reads were joined using fastq-join through the join_paired_ends.py script from QIIME (Caporaso, Kuczynski, et al. 2010) with a minimum overlap of 40 nt (‘--min_overlap40’) and 15% maximum difference between reads (‘--perc_max_diff 15’). Operational taxonomic units (OTUs) were picked using the pick_open_reference_otus.py script in QIIME with the ‘--suppress_step4’ option. (This option skips the final de novo OTU picking step since preliminary analyses indicated it found a very large number of rare OTUs that were subsequently filtered out.) The default phylogenetic comparisons among OTUs were recalculated using Clustal Omega (Sievers et al. 2011) to align sequences because one clade of Methylobacteria consistently misaligned with the default methods, resulting in extremely long branch lengths. The raw OTU table was then filtered to include only samples with at least 10% more reads than the included blanks (ignoring one blank that had obvious non-sample contamination), and to exclude any chloroplast and mitochondria OTUs. This resulted in a BIOM file with 540 samples and 9057 OTUs. All bioinformatic scripts and pipelines used in this analysis are available as part of the associated workflow package via Figshare (DOI: 10.6084/m9.figshare.5886769)

### OTU normalization

OTUs were normalized to 10,000 reads by converting OTU counts to relative abundances, multiplying by 10,000, and rounding the result. (This is essentially taking the average of an infinite number of rarefactions to 10,000.) Any OTUs with counts of 0 across all samples were then filtered out, as were any samples with fewer than 10,000 counts in the original dataset. The nested pie chart of OTU abundance was created with Krona (Ondov, Bergman, and Phillippy 2011) and modified in Inkscape (https://inkscape.org/) to improve clarity.

### Higher-level taxonomy

Counts for higher-level taxonomic clades were constructed from the raw (not normalized) OTU table by summing all read counts within a taxon (species, genus, etc.) based on their uclust-assigned taxonomy. This new OTU table was then normalized to 10,000 reads as above. The read depth for normalization was based on the raw OTU file, not the one with higher-level taxonomy, since the higher-level taxonomy includes each read multiple times (once for OTU, once for species, once for genus, etc.). This resulted in a total of 381 OTUs or higher-level clades across 425 samples.

### Diversity statistics

Alpha and beta diversity were calculated from the normalized OTU table using QIIME. All available alpha diversity metrics were used except for Michaelis-Menten, since this one caused the program to hang. Beta diversity was calculated with weighted UniFrac, unweighted UniFrac, and Bray-Curtis distance metrics, and the first 5 PCs of each used for downstream analysis. This resulted in a total of 49 diversity metrics across 425 samples.

### Predicted Metagenome

The metagenome content of samples was predicted using PICRUST (Langille et al. 2013) according to its recommended protocol. Briefly, the normalized OTU table was filtered to remove non-reference OTUs and then normalized by 16s copy number. Metagenome content was predicted using both COG and KEGG annotations, which were then categorized by function into higher-level groupings. This resulted in 9041 predicted metagenome annotations across 425 samples.

### Maize Genotypes

Maize genotype data was taken from the Maize Diversity Project (http://www.panzea.org). Genotypes consisted of the public Maize GBS v2.7 dataset mapped onto maize genome AGPv3 coordinates. Genotypes for the Goodman Association Panel were subsetted out of the full dataset. When more than one genotype sample was available for an inbred line, the one with the highest coverage was chosen. Genotypes were filtered to remove sites with minor allele frequencies <5% or with fewer than 100 samples with called (=not missing) genotypes. These resulted in a genotype file with 378,686 single-nucleotide polymorphisms (SNPs) across 323 taxa (samples). This genotype file is included in the workflow dataset on Figshare (DOI: 10.6084/m9.figshare.5886769).

### Calculation of Best Linear Unbiased Predictors (BLUPs)

The 425 leaf samples consisted of many plots that were sampled twice (day and night) and a few maize inbred lines that were present at multiple plots in the field. To reduce these to a single value per maize line, best linear unbiased predictors (BLUPs) were calculated using mixed linear models in R (R Core Team 2017). Specifically, the following model was used with the R package lme4 (Bates et al. 2015):

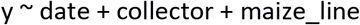

where *y* is each individual phenotype (OTU, diversity metric, metagenome annotation, etc), *date* is the date the sample was collected on, *collector* is the names of the pair of people who collected each sample, and *maize_line* is the name of each maize genotype (B73, Mo17, etc.) All explanatory variables were fit as random effects. Additional cofactors were tested, but all were highly collinear with date and/or collector, and were not significant once these two were included.

Since many phenotypes have highly non-normal distributions, both the raw value and the log-transformed values were fit, and the model with smaller residual error was kept. Other models, including zero-inflated negative binomial and logistic regression models, were tested but resulted in high rates of the generalized linear model failing to converge.

### Narrow-sense heritability analysis

Narrow-sense heritability was calculated for each of the phenotypes using TASSEL (Bradbury et al. 2007). A kinship matrix was calculated from the maize genotype file and fit in a mixed linear model with flowering time as a covariate. (The heritability of flowering time itself was calculated without covariates.) Narrow-sense heritability was estimated as the amount of BLUP phenotypic variance explained by the kinship matrix relative to the total phenotypic variance. Statistical significance of this estimate was determined by randomly permuting the phenotypes and re-calculating heritability to determine a null distribution. For the sake of computation, 100 random permutations were run first to filter out traits with p>0.02, and then 10,000 permutations run on the remaining traits. Empirical p-values were calculated as the fraction of random permutations with heritabilities equal to or greater than the true value.

### Genome-wide association analysis

Genome-wide association for each trait was performed in two steps. First, a mixed linear model (MLM) was fit for each chromosome including both flowering time as a covariate and a kinship matrix constructed from all other chromosomes. (Meaning, for example, that the kinship matrix for chromosome 1 was constructed from markers on chromosomes 2-10; this is the leave-one-out method of Yang et al. (2014).) Since missing values in the flowering time data would have removed ˜10% of samples from the analysis, these missing values were interpolated based on established flowering times at Panzea.org.

After running the initial MLM, empirical p-values were estimated by randomly permuting genotype labels 100 times and re-running the analysis. (Functionally, this is randomizing genotypes among samples while keeping phenotypes and kinship constant; the latter is important to preserve the covariance between phenotypes and kinship.)

GWAS was run on all 255 traits with h^2^≤0.001, consisting of 247 metagenome traits, 5 OTUs or higher-level clades, 2 diversity metrics, and flowering time. Twenty-five metagenome traits were excluded from subsequent analyses both because of their exceptionally high heritabilities (>0.98) and because manual inspection indicated that they were likely artifacts caused by outliers.

### Clustering of GWAS hits

Since several nearby SNPs can be in high linkage, GWAS hits were grouped according to the method of geneSLOPE (Brzyski et al. 2017), picking only the most significant SNP within each linkage window. (Linkage windows were defined by a linkage disequilibrium score of >0.3 as calculated by TASSEL). Clusters of these independent GWAS hits were then identified by dividing the genome into non-overlapping windows of 100,000 base pairs and counting the number of unique traits with a GWAS hit in that region. (“Hits” were defined as a marker with an empirical p-value below the specified threshold, such as 0.05 or 0.01.) Since many traits were correlated with each other, these counts were corrected based on the correlation among traits in each window as follows:

1. The first trait in a window adds a score of 1, representing 1 hit
2. Each additional trait adds a score of (1−r^2^_max_), where r^2^_max_ is the highest correlation between it and any other trait already included in this window. For example, if trait X is the 4^th^ trait counted in a window and has r^2^ values of 0.65, 0.42, and 0.87 with the previous 3 traits, it would add (1 − 0.87) = 0.13 to the score for that window.

### Software

The following software was used as part of this analysis:

- BIOM 2.1.5 (McDonald, Clemente, et al. 2012)
- Clustal Omega 1.2.4 (Sievers et al. 2011)
- FASTX-toolkit 0.0.14 (http://hannonlab.cshl.edu/fastx_toolkit)
- GNU parallel 20130922 (Tange and Others 2011)
- KronaTools 2.7 (Ondov, Bergman, and Phillippy 2011)
- QIIME 1.9.1 (Caporaso, Kuczynski, et al. 2010), which wraps fastq-join (Aronesty 2011), uclust (Edgar 2010), PyNast (Caporaso, Bittinger, et al. 2010), the Geengenes reference alignment (DeSantis et al. 2006; McDonald, Price, et al. 2012), RDP classifier (Wang et al. 2007), FastTree (Price, Dehal, and Arkin 2010), UniFrac (Lozupone and Knight 2005), and Emperor (Vázquez-Baeza et al. 2013)
- Python 3.4.3 and add-on package s argparse 1.1, biokit 0.1.2, biom-format 2.1.5 (McDonald, Clemente, et al. 2012), json 2.0.9, matplotlib 1.5.0 (Hunter 2007), networkx 1.11 (Hagberg, Swart, and S Chult 2008), numpy 1.10.1 (van der Walt, Colbert, and Varoquaux 2011), pandas 0.16.2 (McKinney and Others 2010), re 2.2.1, scipy 0.16.1 (Jones, Oliphant, and Peterson 2016), seaborn 0.8.1, statsmodels 0.6.1. Packages not included in the standard python library were obtained from the python package index (pypi.python.org) using pip.
- R 3.3.3 (R Core Team 2017) and package s argparse 1.0.2 (by Allen Day., n.d.), car 2.1-4 (Fox and Weisberg 2011), colorspace 1.2-6 (Ihaka et al. 2015), corrplot 0.77 (Wei and Simko 2016), gplots 3.0.1 (Warnes et al. 2016), Hotelling 1.0-4 (Curran 2017), KEGGREST 1.12.3 (Tenenbaum 2016), lme4 1.1-12 (Bates et al. 2015), parallel 3.4.0 (R Core Team 2017), and plotrix 3.6-4 (Lemon and Others 2006)
- TASSEL 5.2.26 (Bradbury et al. 2007)

Most analyses were run on a desktop workstation with a 3.5 GHz Intel Xenon E5-1620V3 processor processor (4 physical cores with 2 virtual cores each) and 64 GB of RAM running Linux Mint 17.2. Analyses with large workloads were run on UGA’s Sapelo computing cluster using two 48-core, 256 GB RAM compute nodes. The bioinformatic pipeline for this analysis is available on Github (https://github.com/wallacelab/2014_maize_phyllosphere), and the pipeline plus support files and key intermediate files is available on Figshare (DOI: 10.6084/m9.figshare.5886769).

## Results

### Diversity of the maize leaf microbiome

We sampled the bacterial diversity of approximately 300 diverse maize lines growing in Aurora, New York, in the summer of 2014. Metabolically active bacteria were profiled by extracting total RNA from each sample and using RT-PCR to amplify the V5/V6 region of bacterial 16s rRNA using anti-chloroplast primers (see Methods); a similar focus on RNA instead of DNA was shown to increase power for analyzing the mouse skin microbiomes (Belheouane et al. 2017). Operational Taxonomic Units (OTUs) were clustered with the QIIME pipeline (Caporaso, Kuczynski, et al. 2010) with the default (97%) similarity cutoff. This resulted in a total of 9057 OTUs across 540 samples, with per-sample read depth ranging from 1912 reads to 3.17M (mean 84,142; median 32,302).

Previous results had indicated that phyllsophere communities are generally low-diversity (Bodenhausen, Horton, and Bergelson 2013; Delmotte et al. 2009), and our results confirm this (**Figure 1**). The vast majority of 16s reads map to the Alphaproteobacteria (74%), most of them belonging to just two families: the Sphingomonads (55% of total) and the Methylobacteria (17% of total). The remaining OTUs fall into the Betaproteobacteria (4%), Gammaproteobacteria (6%), Actinobacteria (7%), Bacteroidetes (6%), Deinococcales (0.9%), candidate division TM7 (0.9%), and other groups (1.2%).

**Figure 1.**
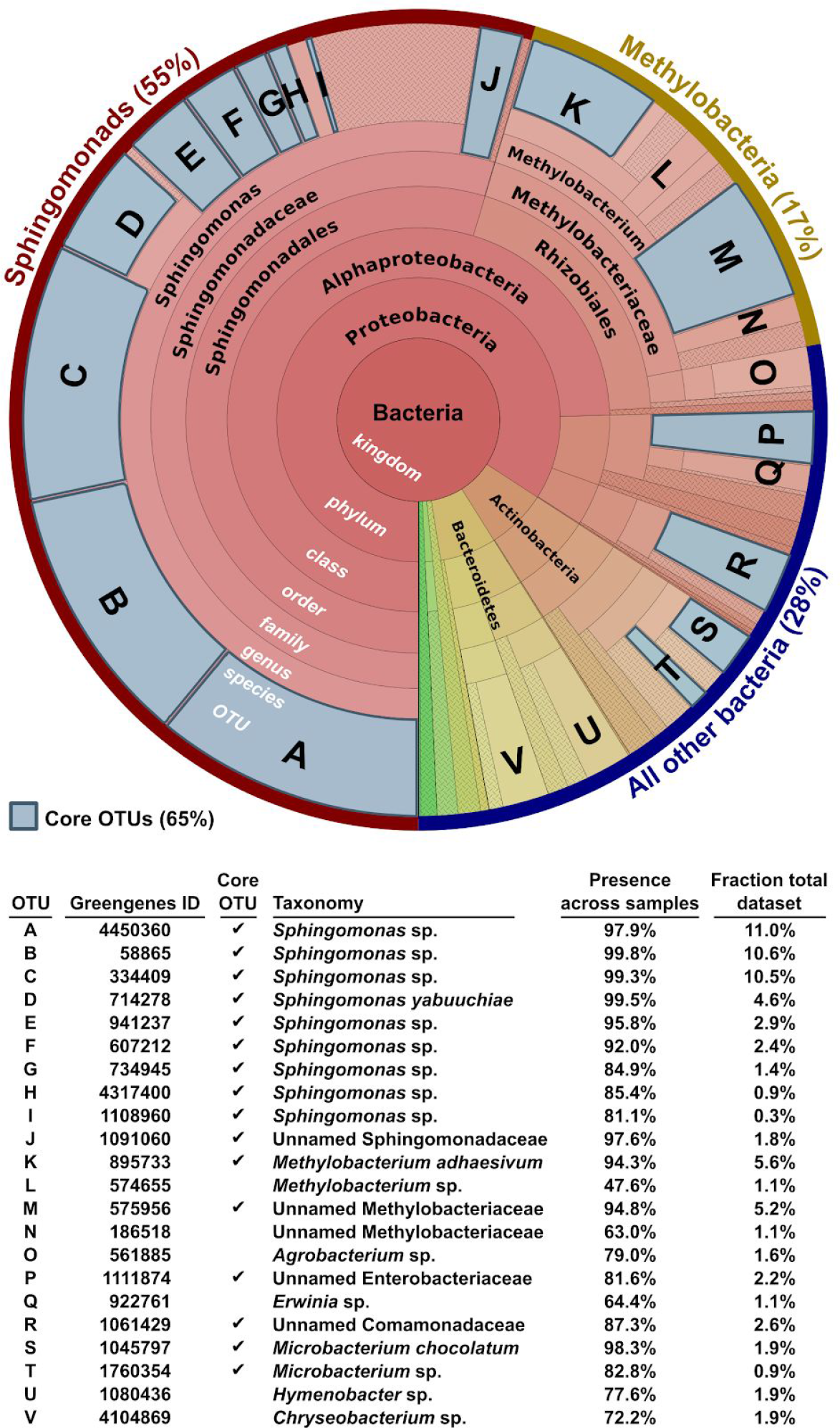
Bacterial diversity of the maize phyllosphere. The distribution of 16s reads among taxa is shown as a Krona plot (Ondov, Bergman, and Phillippy 2011), with the kingdom Bacteria at the center and successively finer taxonomic divisions toward the edge. Core OTUs (present in 80% or more of samples) are shown in blue, and all core OTUs and other abundant OTUs (>1% of total dataset) are labelled with information provided in the table.

We defined the “core” leaf microbiome as those OTUs which were present in at least 80% of samples. Only 16 OTUs fit this category, consisting of 10 Sphingomonads (Alphaproteobacteria; 46.4% of total dataset), 2 Methylobacteria (Alphaproteobacteria; 10.7%), 1 Enterobacteriaceae (Gammaproteobacteria; 2.2%), 1 Comamonadaceae (Betaproteobacteria; 2.6%), and 2 Microbacteria (Actinobacteria; 2.8%). Together these OTUs constitute 64.7% of the total dataset. Other OTUs present at high amounts (>1% of the total dataset) but not in the core community were 2 Methylobacteria (Alphaproteobacteria; 2.2%), a Hymenobacter species (Bacteroidetes; 1.9%), an Erwinia species (Gammaproteobacteria 1.1%), a Chryseobacterium species (Bacteroidetes; 1.9%), and an Agrobacterium species (Alphaproteobacteria; 1.6%) (**Figure 1**). These abundant OTUs make up another 8.6% of the dataset, with the remaining 26.7% of data coming from many (˜8000) OTUs at <1% overall abundance. Some of these low-level OTUs probably derive from sequencing errors or other artifacts, though many of them are probably true individuals present at very low frequency.

The mean fraction of core OTUs in each sample was 64.7% (median 66.1%), with a range from 20.6% to 88.4% (**Figure 2A**). The distribution of major bacterial clades within individual samples followed the same general pattern of the overall dataset; specifically, the majority of sample reads map to Sphingomonads or Methylobacteria, with other clades less common. A few outliers are apparent, however, where normally rare clades make up the majority of reads within a small number of samples. Individual samples show this same pattern (**Figure 2B**), although sample-to-sample variability is still quite strong.

**Figure 2.**
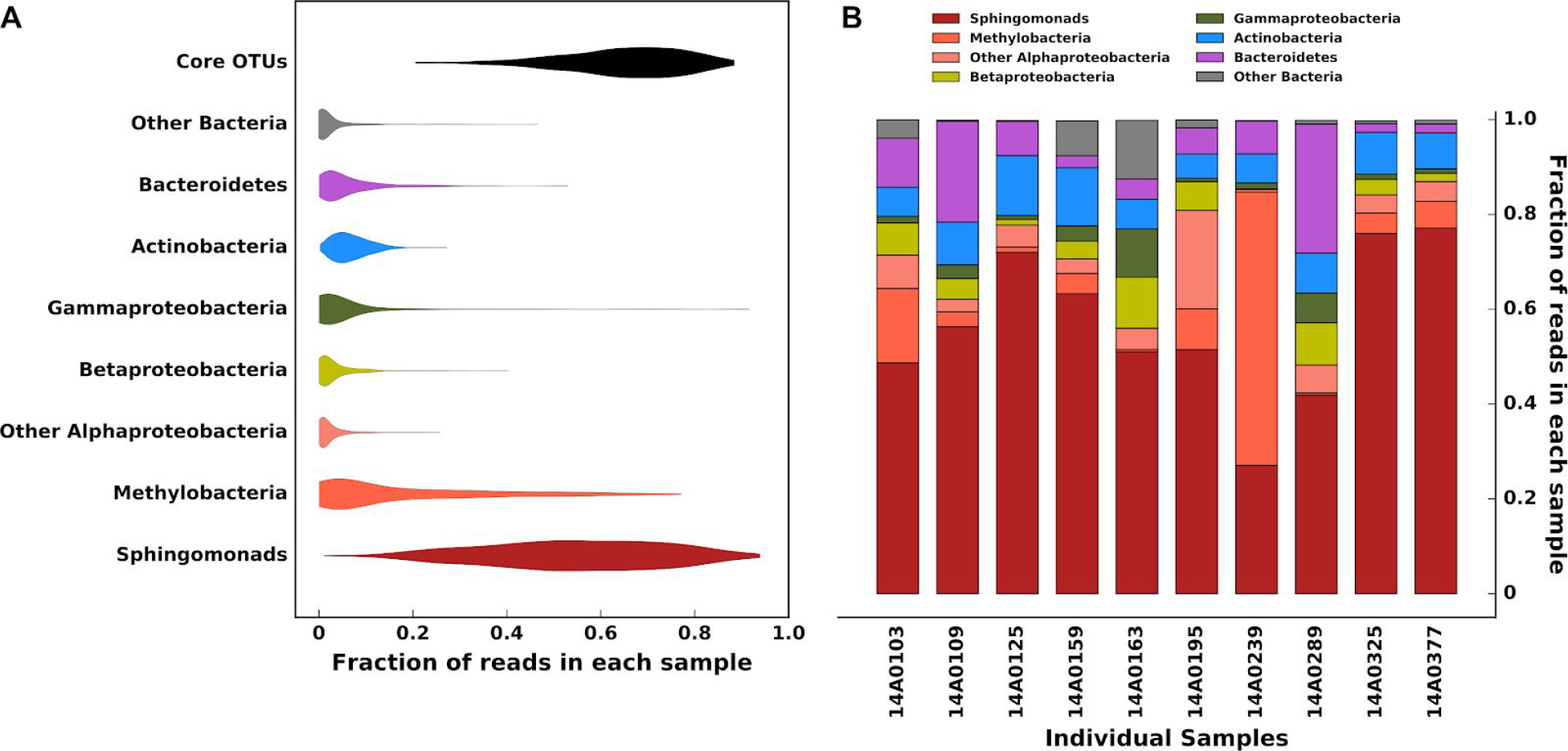
Distribution of major taxonomic divisions in samples. (A) The distribution of different groups of taxa are shown according to their fraction of total reads within each sample. Although most samples are dominated by Spingomonads, Methylobacteria, and other Core OTUs, outliers are apparent based on the presence of small numbers of samples with extremely large values for some taxa. (B) The distribution of these same taxa (except for Core OTUs) is shown in a random subset of individual samples. The colors are the same as in (A), and significant sample-to-sample variability is apparent.

### Sample alpha diversity

We performed several analyses of the bacterial alpha diversity, or diversity present within individual samples. There are many different ways to measure alpha diversity, so rather than focus on any individual measure we looked at patterns across them. We first calculated alpha diversity using all metrics available in QIIME except for Michaelis-Menten’s fit (because it hung the program without ever terminating); the Kempton-Taylor Q metric was later dropped because it generated only NA results. Many of the available metrics measure highly similar properties of the data, so we pruned them to an informative subset where no pair of metrics had a correlation ≥ 0.9 or ≤ −0.9.) (**Supplemental Figure 2**). This left seven metrics: doubletons, Strong’s index, Berger-Parker dominance, Heip’s evenness, Simpson’s evenness, Chao1 upper bound, and whole-tree phylogenetic distance.

We tested several cofactors for potential influence on these alpha diversity metrics: the date and time of sampling, the collection team, the specific plate the RNA was stored in (for batch effects), and the maize subpopulation (tropical, stiff-stalk, non-stiff-stalk, popcorn, sweet corn, or mixed). The cofactors explained between 0% and 10% of variance individually and between 4% and 15% of variance when combined (**Supplemental Figure 3**). Hidden factor analysis (analogous to principal components analysis; (Lawley and Maxwell 1962)) identified hidden factors that could explain 0-5% more variance, depending on the trait (**Supplemental Figure 4**). A Type II sum-of-square analysis was performed to identify individual cofactors that underlie these patterns. Only one cofactor was significantly associated with a metric after Bonferroni correction (**Supplemental Table 1**): the collection team has a Bonferroni-corrected p-value of 2.1 × 10^-5^ for whole-tree phylogenetic distance. Rarefaction curves of alpha diversity support higher diversity levels for two specific teams, but also for samples from tropical maize lines and plants sampled later in the season (**Supplemental Figure 5**). Due to the nature of the sampling scheme these factors are highly confounded in the data, so we cannot say which of them are actually driving this pattern.

### Sample beta diversity

To look at diversity across samples, we performed beta diversity analysis as implemented in QIIME. Beta diversity was calculated by three metrics: unweighted UniFrac, which calculates distances based on phylogenetic dissimilarity among OTUs (Lozupone and Knight 2005); weighted UniFrac, which does the same but also weights the measures by OTU abundance (Lozupone and Knight 2005); and Bray-Curtis Dissimilarity, an ecological measure comparing species counts in two samples (Bray and Curtis 1957).

We compared the principal coordinates of these three datasets to possible confounding factors to identify technical artifacts that could be driving the apparent community structure (**Supplemental Figure 6**). Neither Bray-Curtis nor Weighted UniFrac show any clear association with batch effects. The unweighted Unifrac measure, however, shows samples clearly dividing into two groups corresponding to the two batches samples were processed in, which are also strongly but not completely correlated with day versus night samples. Unweighted Unifrac is more sensitive to rare OTUs than the other metrics, and the lack of similar clustering for Weighted UniFrac and Bray-Curtis dissimilarity implies that the more common OTUs were not affected by batch processing. MANOVA analysis of the first 10 principal components confirms that the RT-PCR batch has the strongest effect on Unweighted UniFrac PCs, whereas both Bray-Curtis and Weighted Unifrac were most strongly correlated with the collection team (**Supplemental Table 2**) Although many cofactors show some level of statistically significant association with beta diversity metrics (**Supplemental Table 2**), visual inspection shows little practical clustering for most of them (e.g., **Supplemental Figure 6**).

Only the weighted UniFrac measure showed significant correlation with bacterial clades (**Supplemental Table 3**). Correlation values for the most likely clades driving the first 5 principal coordinates are shown in **Table 1**, and they are graphed for the first 3 PCs in **Figure 3**. In several cases multiple taxonomic levels are highly correlated with a single PC, usually because higher-level clades consist almost entirely of reads from a single lower-level clade. Not surprisingly, the two most abundant clades--Methylobacteria and Sphingomoinads--have a large role in driving community structure. PC1 appears to be driven by the family Methylobacteriaceae (r^2^=0.845), while PC3 is driven by the Sphingomonadaceae (r^2^=0.552). (Although PC3 is slightly more correlated with the Gammaproteobacteria--r^2^=0.571--it seems more likely that the much more abundant sphingomonads are actually driving that PC). Surprisingly, PC2 is driven by the Bacteroidetes (r^2^=0.602), a relatively low-abundance clade that comprises only 6% of the overall dataset. It does, however, contain the two most-abundant non-core OTUs in the dataset (**Figure 1**), though one of these (Hymenobacter) also appears to be the major driver of PC4 (r^2^=0.699). Finally, PC5 appears to be driven by the Actinobacteria (r^2^=0.609), the other major phylum in the data and which contains two core OTUs (**Figure 1**).

**Table 1.** **Bacterial clades likely driving beta diversity**

| PC #<sup>a</sup> | Clade | r<sup>2</sup> |
| --- | --- | --- |
| 1 | Methylobacteriaceae (family) | 0.845 |
| 2 | Bacteroidetes (phylum) | 0.602 |
| 3 | Sphingomonadaceae (family) | 0.552 |
| 4 | Hymenobacter (genus) | 0.699 |
| 5 | Actinobacteria (phylum) | 0.609 |
| *a - All PCs are from weighted UniFrac analysis* |  |  |

**Figure 3.**
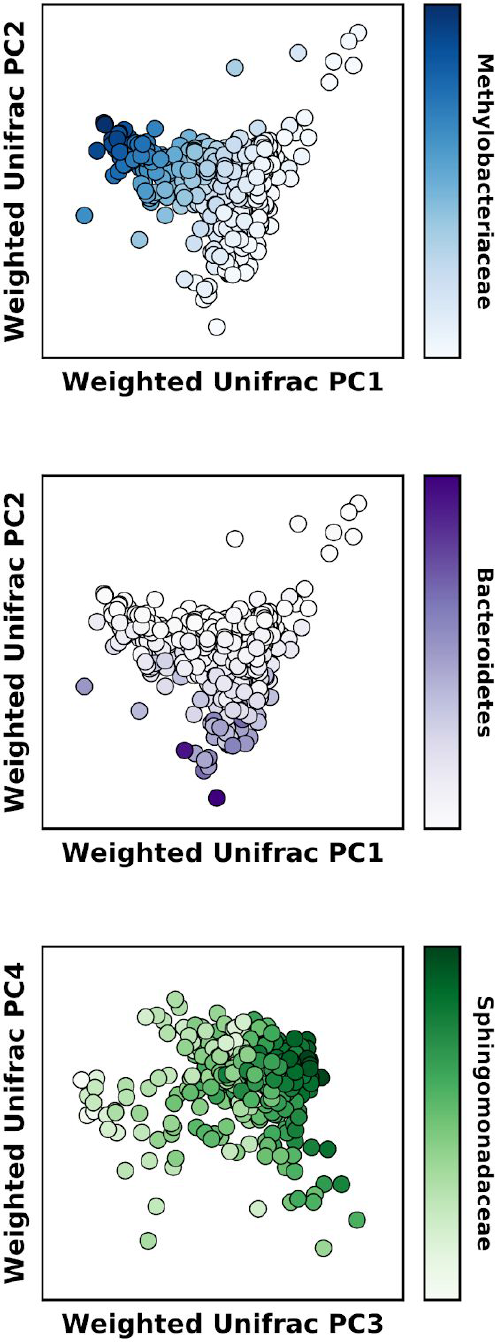
Major drivers of phyllosphere beta diversity. The first several principal coordinates of Weighted UniFrac beta diversity were compared to the taxonomic clades present in the data to identify which taxa may be driving the overall community makeup. PC1 is most strongly correlated with the Methylobacteriaceae and PC2 with Bacteroidetes. Although PC2 has a slightly stronger correlation with Gammaproteobacteria than the Sphingomonadaceae (**Supplemental Table 3**), the much greater abundances of Sphingomonads (**Figure 1**) implies that this clade is actually driving the PC.

### Consistency within field plots

Prior research has shown that individual phyllosphere species can have patchy distributions (Steven E. Lindow and Brandl 2003; Monier and Lindow 2004). It was unclear whether our sampling scheme would capture enough leaf surface to be representative of the entire field plot. To determine this, we compared paired day-night samples within each plot other plots in the experiment. Within-plot samples are visibly more similar to each other than to neighboring plots (**Figure 4**). We further analyzed the distance between samples through various beta-diversity metrics. These show that within-plot sample pairs are significantly (p <= 0.01) more similar to each other than they are to corresponding samples from neighboring plots, to any non-neighboring plot, or to an equivalently sized random subset of non-neighboring plots. The only exception was among samples taken on 26 Aug and measured with weighted UniFrac distance, in which case the within-plot and neighboring-plot distances were not statistically different (**Supplemental Figure 7**). The distances to neighboring-plot samples were significantly smaller than distances to random non-neighbor plots in 5 out of 9 comparisons, indicating there may be some spatial autocorrelation within the field samples.

**Figure 4.**
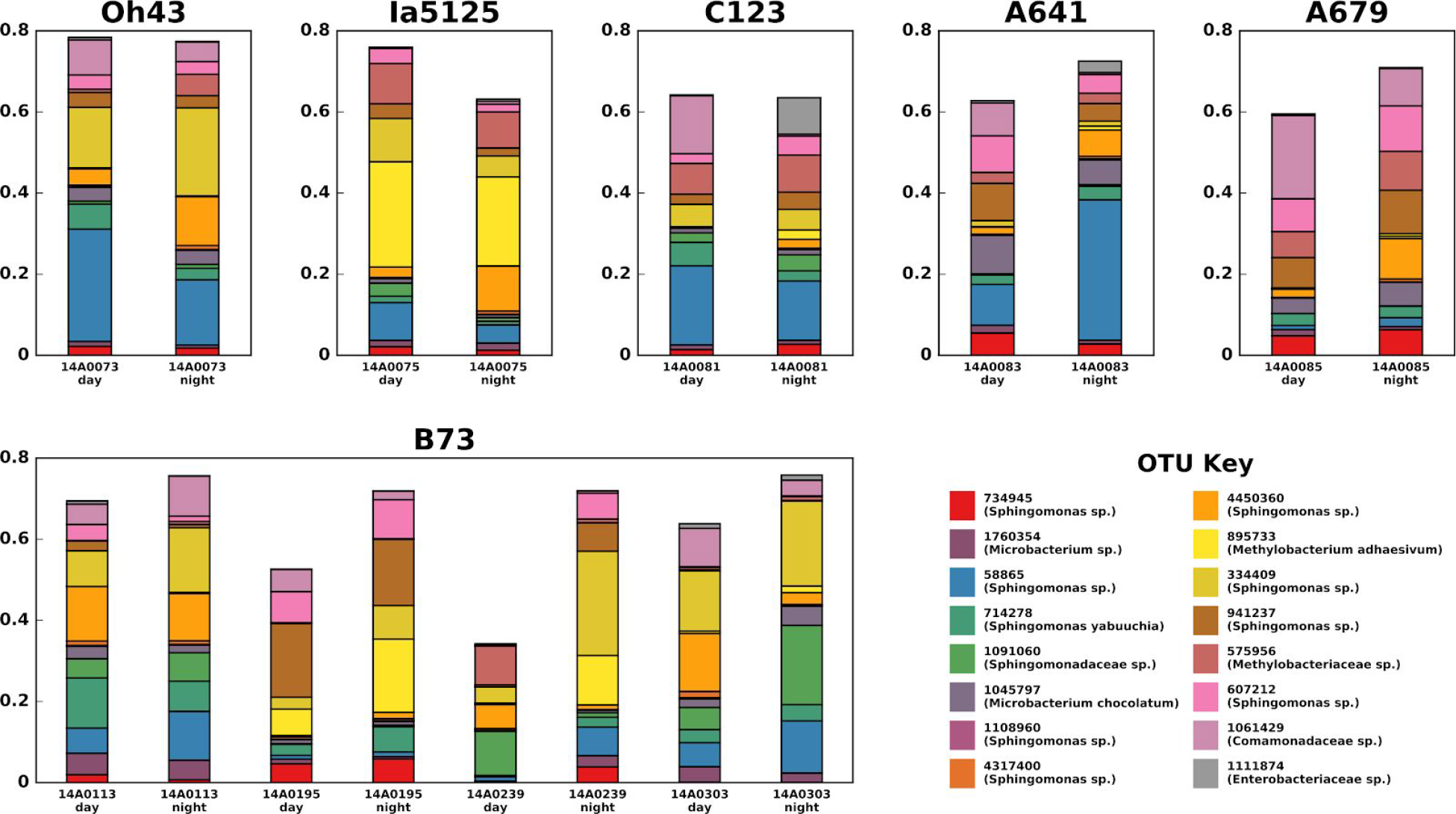
Comparison of paired samples. (Top) Five neighboring plots are shown with the distribution of core OTUs in their paired day and night samples. Variation between day and night is small relative to variation from plot to plot. (Bottom) Paired day-night samples for four plots of one genotype, B73, are shown. Although day-night samplings are relatively consistent (except for 14A0239), there is no more apparent similarity among B73 samples than among neighboring samples with different genotypes.

### Heritability of microbiome community characteristics

To estimate how much maize genetics shapes its overall phyllosphere community, we estimated the heritability of alpha and beta diversity metrics using the Best Linear Unbiased Predictors from the data. (Some data was log-transformed due to giving a better fit; see Methods.) Heritability in this sense does not refer to vertical transmission but rather to how much of total phenotypic variation is due to plant genetics. In essence, we are treating characteristics of the phyllosphere microbiome as a plant phenotype and estimating how much control the plant itself exerts over the trait. When testing the overall microbial community instead of individual microbes, this is the same as the community heritability (H^2^_c_) advocated by van Opstal and Bordenstein (van Opstal and Bordenstein 2015), although our methods estimate narrow-sense heritability (h^2^, the proportion due to additive genetic effects) instead of broad-sense heritability (H^2^, the proportion due to total genetics).

We tested all alpha diversity metrics available in QIIME, along with the first 5 principal coordinates from Bray-Curtis, Unweighted UniFrac, and Weighted UniFrac beta diversity analyses. Each metric was individually fit as the response variable in a mixed linear model in TASSEL (Bradbury et al. 2007). Narrow-sense heritability (h^2^) was estimated by including a kinship matrix generated from public genotype data on these maize lines and estimating the proportion of total variance explained by the kinship matrix. Alpha diversity metrics were extremely poorly heritable, with values ranging from 0 (most traits) to 0.05 (log-transformed Strong’s dominance index) (**Supplemental Table 4**). Beta diversity was significantly more heritable, with maximum heritability of 0.598 for Weighted Unifrac PC1, 0.521 for Unweighted UniFrac PC2, and 0.19 for Bray-Curtis PC1 (**Supplemental Table 5**).

To determine the significance of these heritability scores, we performed a permutation analysis where the phenotype data was randomly shuffled relative to the kinship matrix and the heritability recalculated. We initially screened all metrics against 100 random permutations; metrics with an empirical p-value ≤ 0.02 were then rerun against 10,000 random permutations to more precisely estimate the null distribution. Metrics with a final empirical p-value ≤ 0.001 were considered significant. (That is, at most 1 random permutation in 1000 had a heritability as high as or higher than the actual data.) None of the alpha diversity metrics passed the initial filter, while only two beta diversity metrics passed the second: Weighted Unifrac PC1 (h^2^=0.598, p = 0.0003), and Unweighted Unifrac PC2 (h^2^=0.521, p = 0.0003).

### Heritability of bacterial OTUs and clades

Using the same methods as for alpha and beta diversity, we performed heritability analysis on 185 individual OTUs and 196 higher taxonomic units (species, genus, etc.). We tested several methods to deal with the non-normality of many OTU counts, including log transformation and several variations on negative binomial regression. In the end, we chose log-plus-1 transformation of normalized OTU counts because it was the most robust across OTUs. (The various negative binomial variants had a high tendency to fail due to the generalized linear model not converging.)

Most taxa are not significantly heritable, with only 2 OTUs and 3 higher-level taxonomic clades showing significant heritability at an empirical p-value ≤0.001 (**Figure 5**). All five of these are in the class Alphaproteobacteria; four of the five are in the order Rhizobiales, and three are in the family Methylobacteria. An additional 376 OTUs or higher-level clades did not show significant heritability, indicating that the majority of taxa living on the maize leaf surface are minimally influenced by the genotype of their host.

**Figure 5.**
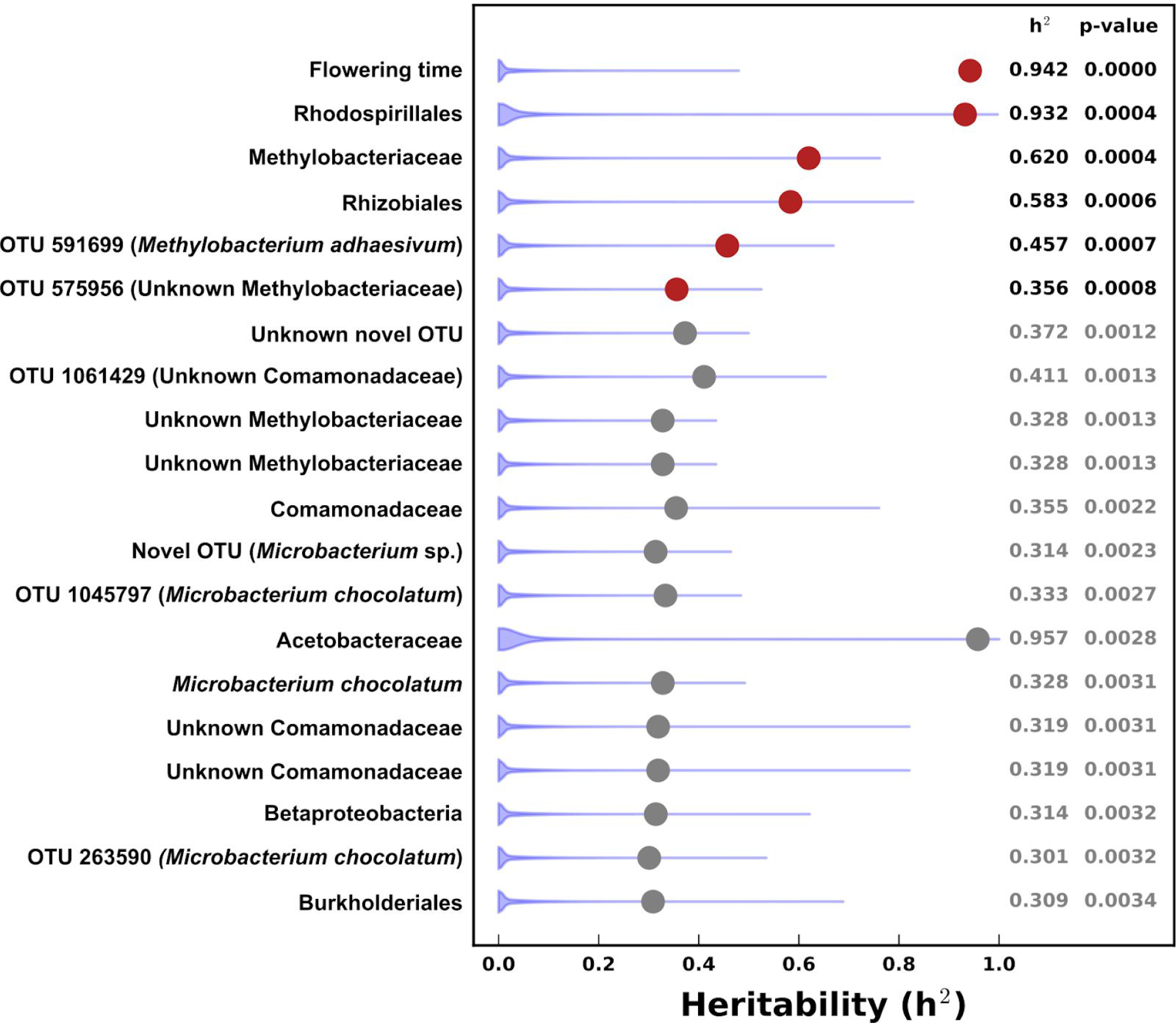
Heritability of OTUs. Narrow-sense heritability (h^2^) was calculated using a mixed linear model for 185 individual OTUs and all 196 of their higher-level clades, plus flowering time as a positive control. Each heritability value was then compared to that obtained from 10,000 random permutations of the data to yield empirical p-values. Traits with empirical p-values ≤ 0.001 were considered significantly heritable (red); all others were considered not significant (gray).

Of the 16 core OTUs, only one showed significant heritability (h^2^ = 0.356, p = 0.0008): OTU 575956, an unnamed Methylobacterium. The heritability of other core OTUs ranged from 0 to 0.411, and only two others come close to statistical significance: an unnamed Comamonadaceae at h^2^=0.411, p = 0.0013, and *Microbacterium chocolatum* OTU 1045797 at h^2^ = 0.333 and p=0.0027. (**Supplemental Table 6**).

### Heritability of inferred metagenome content

It is increasingly recognized that the functional capacity of a microbiome may be both more important and more consistent than its taxonomic makeup (Human Microbiome Project Consortium 2012). We used PICRUST (Langille et al. 2013) to infer the metagenome content of the maize leaf microbiome using annotations from both the Kyoto Encyclopedia of Genes and Genomes (KEGG Orthology, KO) (Kanehisa and Goto 2000) and Clusters of Orthologous Genes (COG) (Tatusov et al. 2000). We then calculated the heritability of the inferred metabolic annotations as above, including binning individual metabolic annotations into groups according to their respective hierarchies.

The raw number of metagenome annotations (9041) was much larger than either the diversity metrics (48) or the OTUs and taxonomy (381). A correspondingly larger number of metagenome annotations were identified as significantly heritable (empirical p-value ≤0.001), with 120 KO terms and 127 COG terms (**Supplemental Table 7**). Twenty-five of these terms (11 KO terms and 14 COG terms) appeared to be artifacts due to highly skewed distributions (**Supplemental Figure 8**) and were excluded from further analysis.

The significantly heritable terms spanned a large range of metabolic and cellular functions (**Figure 6**, **Supplemental Figure 9**). To identify any functions that were enriched in the phyllosphere microbiome, we used the hierarchy of KO or COG terms embedded in the PICRUST-annotated file to search for enriched terms. Parent terms were tested by both Fisher Exact test and by Annotation Enrichment Analysis (AEA) (Glass and Girvan 2014), which uses permutation of the hierarchy to correct for highly connected terms being more likely to be found significant by chance. After correcting for false discovery rate using the Benjamini-Hochberg method (Benjamini and Hochberg 1995), only 4 KO terms and zero COG terms were significantly enriched by Fisher Exact test. (**Table 2**). No terms were significantly enriched by AEA, although the rank-order of p-values was similar (**Supplemental Table 8**).

**Figure 6.**
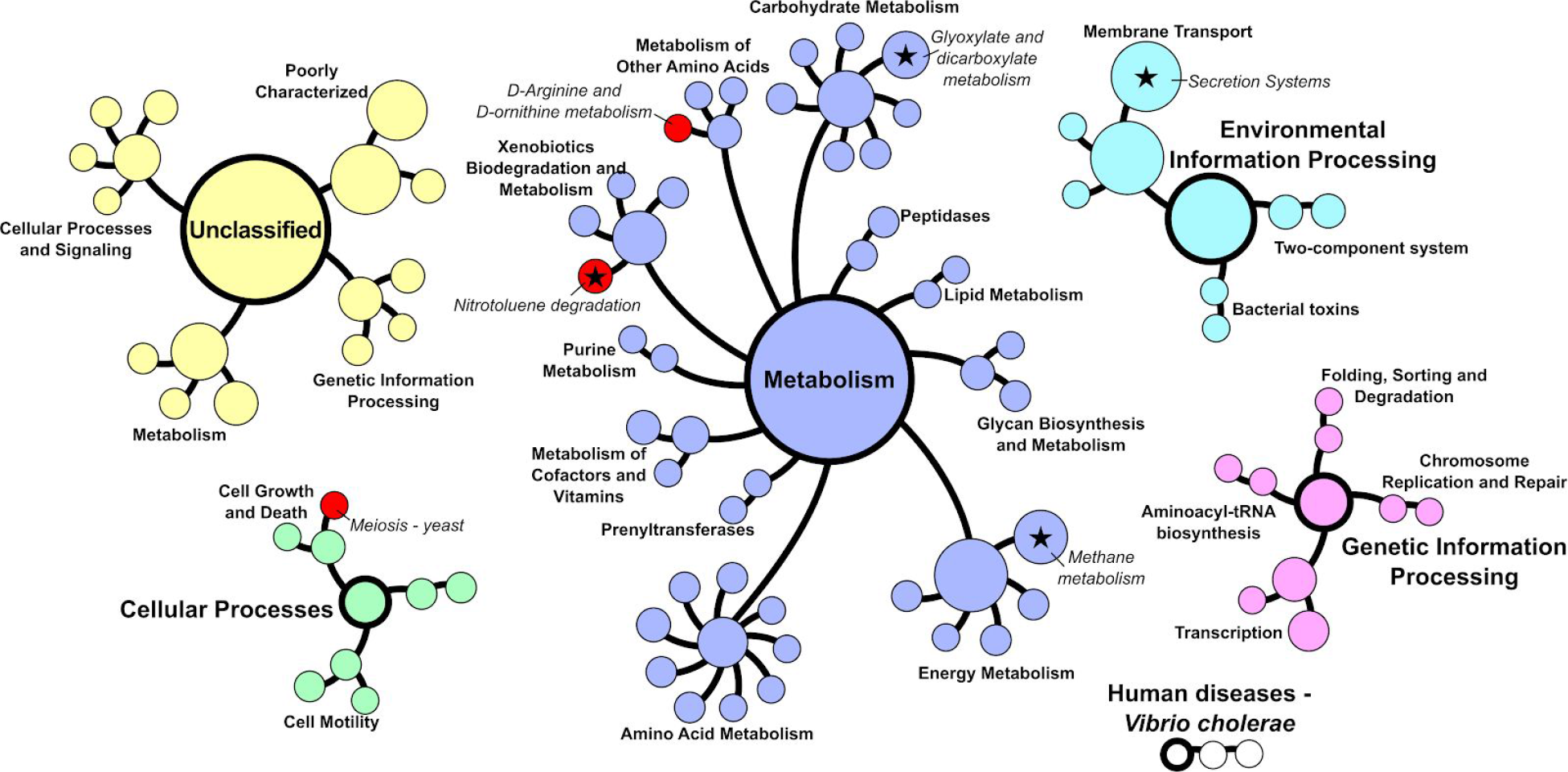
Heritability of inferred metagenome content. Metagenome functional capacity was estimated for each sample by PICRUST (Langille et al. 2013) and analyzed for narrow-sense heritability as per **Figure 5**. The hierarchy of heritable KEGG Orthology (KO) terms is shown, colored by major groupings; individual KO terms are omitted for clarity. (See **Supplemental Figure 9** for COG terms.) Three functional terms that were themselves significantly heritable are shown in red, while stars indicate the four terms that were significantly enriched based on FDR-corrected Fisher’s Exact Test (q ≤ 0.05). Only one term (nitrotoluene degradation) was significant in both analyses.

**Table 2.** **Significantly enriched metagenome terms**

| Term | Fisher Exact FDR<sup>a</sup> | AEA empirical p-value<sup>b</sup> | Individual h<sup>2c</sup> | h<sup>2</sup> empirical p-value<sup>d</sup> |
| --- | --- | --- | --- | --- |
| Methane metabolism | 0.011 | 0.17176 | 0.201 | 0.06 |
| Nitrotoluene degradation | 0.011 | 0.21828 | 0.677 | < 0.0001 |
| Glyoxylate and dicarboxylate metabolism | 0.030 | 0.12159 | 0.137 | 0.07 |
| Secretion system | 0.041 | 0.15167 | 0.479 | 0.003 |
| *a – Fisher's Exact test for enrichment after Benjamini-Hochberg (false discovery rate) correction*<br>*b – Empirical p-value from 100,000 random permutations with Annotation Enrichment Analysis*<br>*c – Estimated narrow-sense heritability of the individual term*<br>*d - Empirical p-value of the heritability estimate based on either 100 or 10,000 random permutations* |  |  |  |  |

### Genome-wide association

To identify specific genes or regions associated with each trait, we performed genome-wide association (GWAS) analysis for all traits with an empirical heritability p-value ≤0.001 (n=230 traits total, including 2 OTUs, 3 higher-order taxa, 2 diversity metrics, 222 metagenome predictions, and flowering time as a positive control). For each trait, a mixed linear model was run separately for each chromosomes using a kinship matrix constructed from all other chromosomes (Lippert et al. 2011; J. Yang et al. 2014). Flowering time was included as a covariate for all traits except itself, with missing flowering times (˜10% of samples) interpolated based on published data for this panel available at Panzea.org. This same analysis was then run with 100 random permutations of genotypes (keeping phenotypes and kinship constant) to estimate empirical p-values for all associations.

The results for flowering time and two other traits (log COG0181 and log K02769) are shown in **Figure 7**. Flowering time associations hit several genes known to be important for this trait from previous studies (Buckler et al. 2009; Dong et al. 2012). Flowering time also has many more hits than the phyllosphere traits, with 10 loci having empirical p-values ≤0.01 compared to only 2 loci for the best phyllosphere traits. Most phyllosphere traits (177 out of 229) do not have any significant hits at p≤0.01 (**Supplemental Table 9**); 49 traits have 1 hit and 3 have 2 hits. Even if the threshold is relaxed to p≤0.05 (a level likely to introduce many false positives), only 136 out of the 229 phyllosphere traits have at least 1 hit.

**Figure 7.**
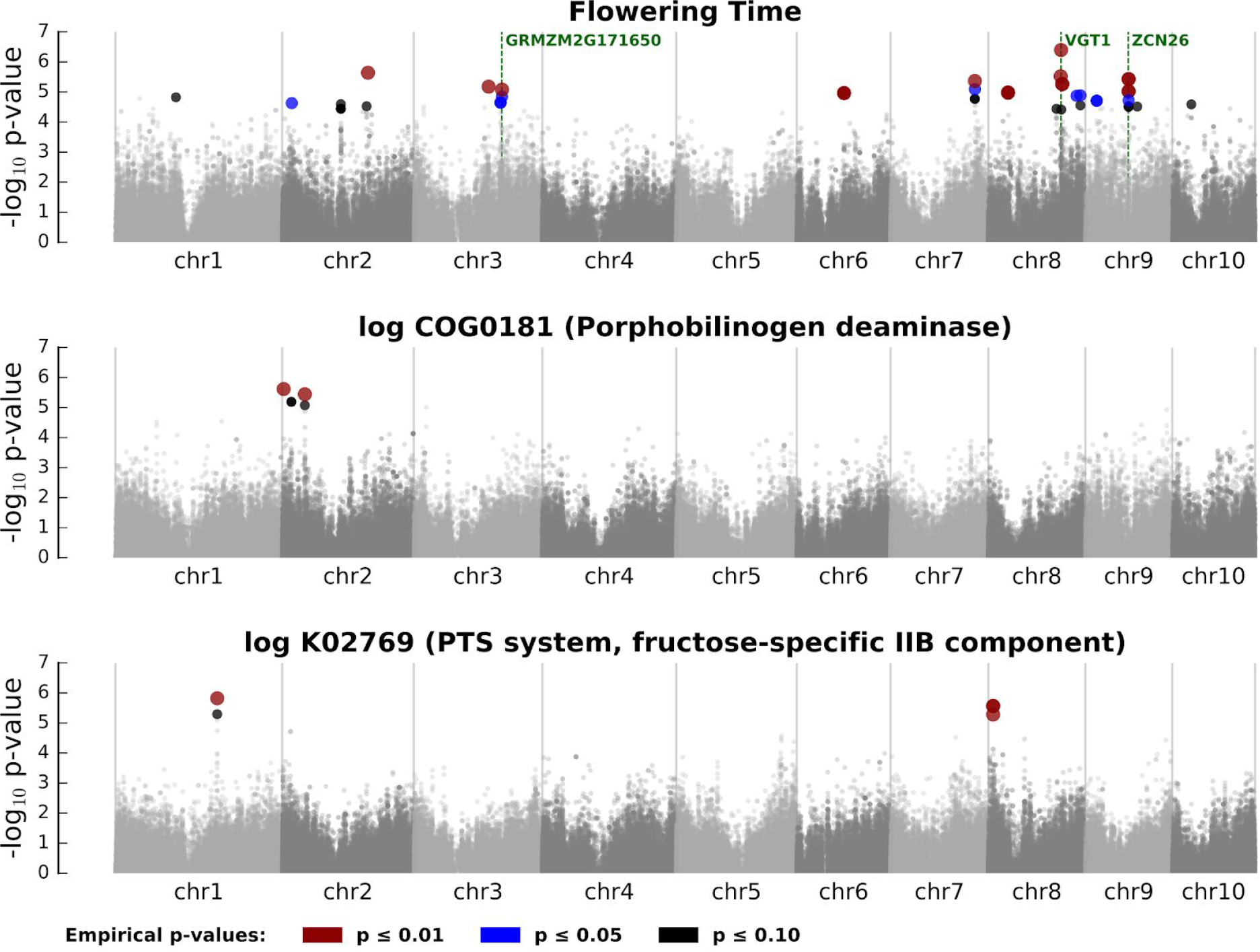
Genome-wide association results. All traits with significant heritability (see **Figures 5 & 6**) were analyzed by genome-wide association using a mixed linear model in TASSEL (Bradbury et al. 2007). The raw results were compared to 100 permuted analyses where genotype labels were scrambled while keeping phenotypes and kinship constant to determine empirical p-values for each hit. The results from the three strongest traits are shown, with chromosome position along the X axis and −log_10_ of the raw p-value on the Y axis. Hits are colored according to their empirical p-value; we consider only hits with an empirical p-value ≤ 0.01 to be statistically significant, though hits with p up to 0.10 are shown. Flowering time (top) shows multiple significant hits, including several near genes known to be involved in flowering. All phyllosphere traits show fewer robust associations, with the two shown (COG0181 and K02769, both predicted metagenome terms) each having only two significant two loci. The hit on chromosome 8 for K02769 is near GRMZM2G396541, a sulfated surface glycoprotein also implicated in Methylobacteria associations (albeit outside out significance cutoff; see main text). All other genes close to these hits have no known function. Most other phyllosphere traits show at most one significant hit, with the majority having none at p ≤ 0.01.

We also investigated the possibility that multiple traits are controlled by the same locus. For this, we separated the genome into 100 kb windows and counted the number of traits with an association in each window. We then adjusted this score based on the correlations among traits (see Methods for details) to generate a correlation-corrected score for each window. We identified two windows with correlation-corrected scores >2 (roughly equivalent to at least two wholly independent traits), on chromosomes 7 and 10, and both result from a large number of traits having significant associations at a single SNP marker (21 traits at 7:165,427,159 and 6 traits at 10:92,038,994). All but one of the traits at these two locations are for metagenome predictions (**Table 3**); the cluster on chromosome 7 also includes a hit for a beta diversity metric, namely, the first PC of weighted UniFrac analysis.

**Table 3.**
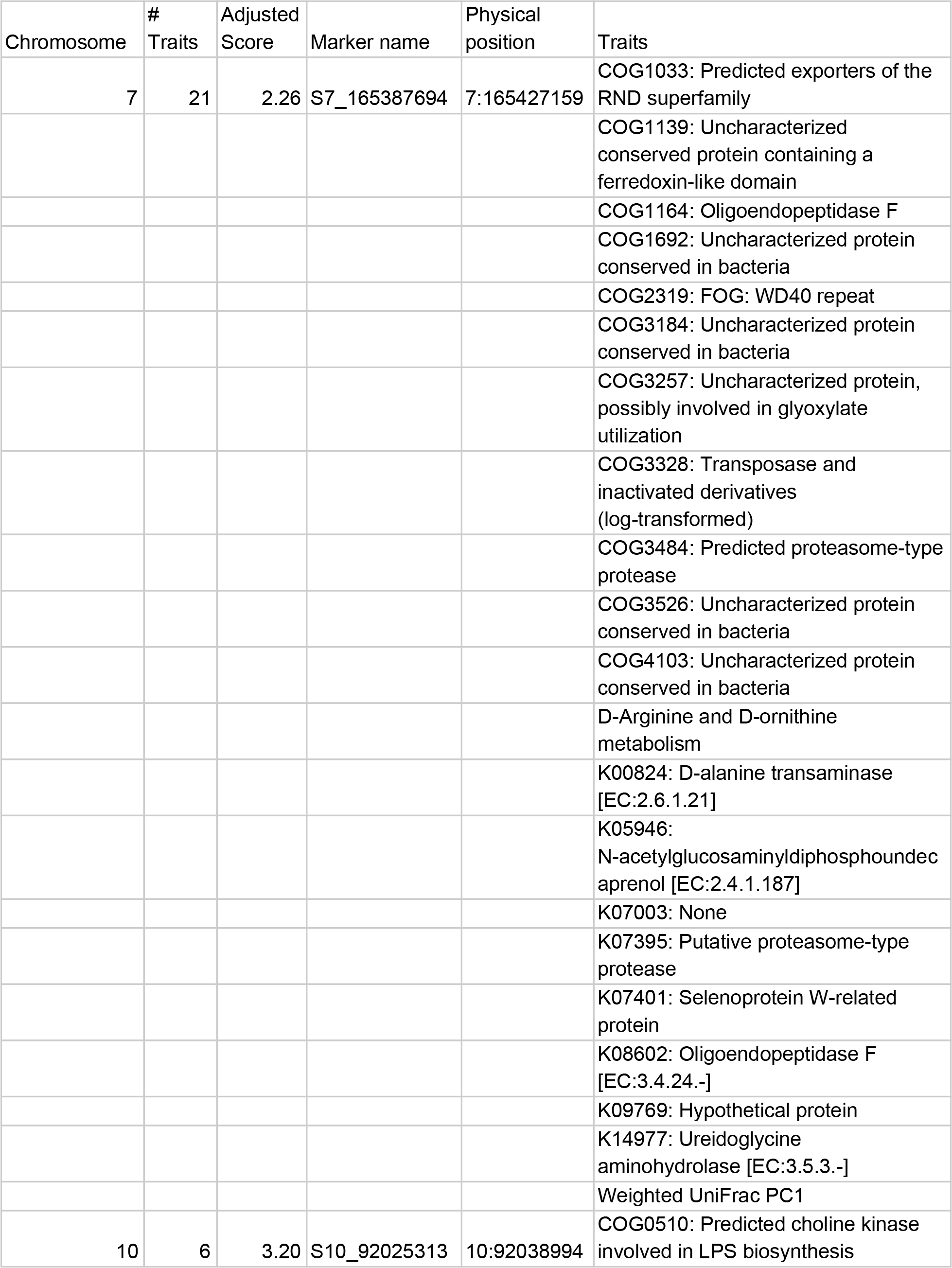
**GWAS hit clusters**

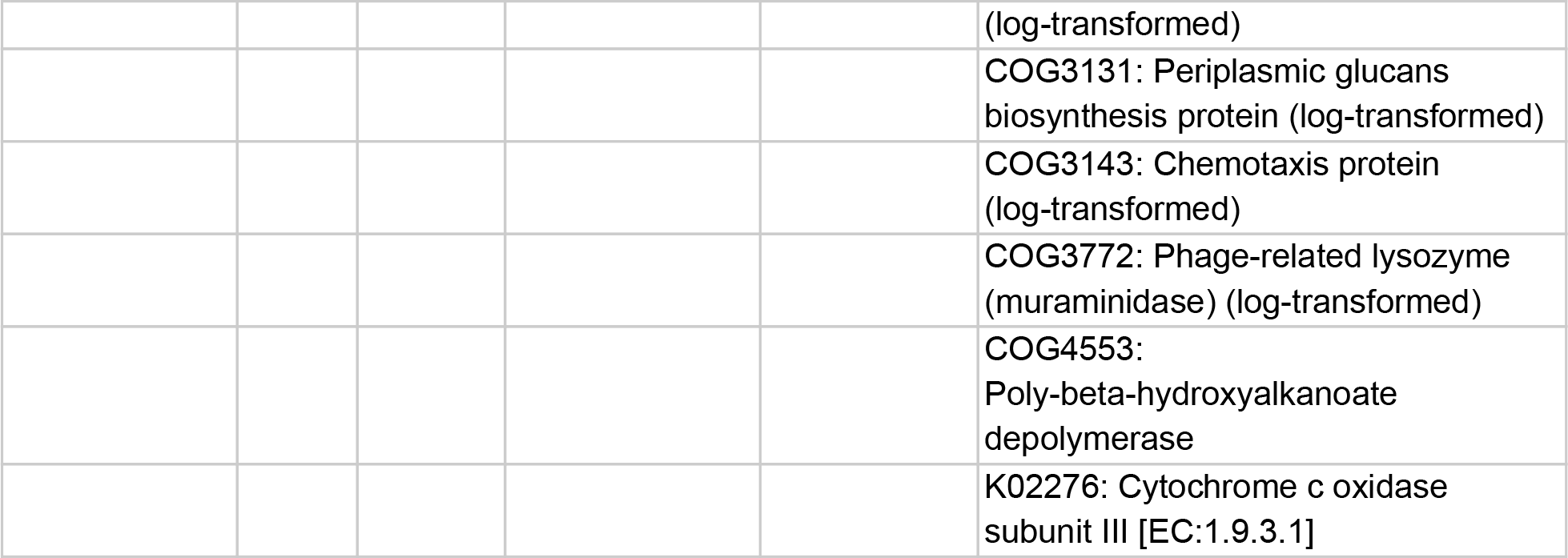

## Discussion

### The maize phyllosphere is low diversity and distinct from the rhizosphere

Our results confirm that the maize phyllosphere is a low diversity community. Over 70% of sample reads map to just two families, the Sphingomonads and the Methylobacteria, with over half (52%) of all reads mapping to a single genus, *Sphingomonas*. These results are in line with phyllosphere data from other species (Delmotte et al. 2009) and in stark contrast with data from the maize rhizosphere, where the overall community shows both more total OTUs and greater taxonomic diversity (Peiffer et al. 2013; Y. Yang et al. 2017). For example, a study that had previously looked at maize rhizosphere samples at this same experimental station showed that the Sphingomonads and Methylobactereia make up <1% of the rhizosphere (Peiffer et al. 2013). This is in line with other studies that show little correlation between soil and leaf microbial communities, especially later in the season (such as when these samples were taken) (Kim et al. 2012; Copeland et al. 2015). Wind dispersal from other plants appears to play a strong role in phyllosphere bacterial community dynamics (S. E. Lindow and Andersen 1996; Lymperopoulou, Adams, and Lindow 2016), so that the local soil may not even serve as the inoculum for much of the phyllosphere community.

### Drivers of phyllosphere community structure

One of the major goals of this research was to identify the primary drivers of the maize phyllosphere community. The first task was to rule out any confounding from technical errors. Both Bray-Curtis and Weighted UniFrac measures of community diversity show significant associations with possible confounding factors, especially the collection team. It is possible that this reflects operational biases for each team. It is also possible, however, that the “collection team” variable is instead just the best representation of some latent variable in the data because it breaks the data into the most number of groups (13), and these groups tend to be confounded with cofactors like flowering time and field location due to the logistics of sample collection. These possibilities are not mutually exclusive, and with the current data we cannot wholly tell which is correct. Regardless of what process it represents, the collection team was included as a cofactor in the generation of best linear unbiased predictors (BLUPs) to factor it out of downstream analyses.

**Figure 4** shows that our paired day and night samples show relatively strong consistency within a field plot, indicating that our sampling method was sufficient to get a good representation of the plant phyllosphere (specifically, that of upper leaves). The visible differences between neighboring plots of different genotypes (**Figure 4, top**) would indicate a potentially strong effect of plant genotype. However, the B73 check line, which was planted at multiple locations throughout the field, shows no more plot-to-plot consistency than neighboring plots with different genotypes (**Figure 4, bottom**). This indicates that there is probably a strong environmental and/or stochastic component to phyllosphere community composition. Since all plants in this experiment grew in the same field and within a few dozen meters of each other, we expect the environmental variation from plot to plot to be minimal, so much of the non-genetic variation in this data likely comes from stochastic noise. This is in line with previous results showing that phyllosphere communities are shaped by stochastic events early in colonization (Maignien et al. 2014).

No alpha diversity metrics show any significant heritability, indicating that maize genotype does not have a major effect on the diversity of its phyllosphere community. This is surprising, as previous work in a maize recombinant inbred line (RIL) population showed a significant relationship between plant genetics, community diversity, and disease resistance (Balint-Kurti et al. 2010). Specifically, that analysis mapped loci influencing Simpson diversity to six bins in the maize genome. In contrast, the Simpson’s diversity index of our data has a heritability of only 0.008 and an empirical p-value of 0.38, so it was never included in the GWAS mapping analysis. (Although this heritability is very low, it is actually among the highest for alpha-diversity metrics.) Many experimental details are different between the two analyses (RILs versus a diversity panel, looking at terminal restriction fragment length polymorphisms versus 16s rRNA, etc.), so we cannot say which is more likely to be correct overall.

Most beta diversity measures were also not significantly heritable, with the exception of UniFrac principal coordinates. Weighted UniFrac PC1 showed significant heritability (h^2^=0.598) and is driven primarily by Methylobacteriaceae (**Figure 3**), a bacterial clade which is also significantly heritable on its own (**Figure 5**). This indicates that plant genetics can significantly impact the degree to which these organisms colonize the phyllosphere, although they were one of very few clades showing significant heritability. Methylobacteria are thought to subsist off of short-chain carbon molecules that leak out of leaves (Steven E. Lindow and Brandl 2003; Vorholt 2012). The association between Methylobacteria and plant genetics may thus relate to different metabolic pathways within the plants that generate these compounds, or to different cell wall structures that make leaves more or less permeable to them. The only other diversity metric with significant heritability was unweighted UniFrac PC2. Although it is only weakly associated with specific bacterial clades, its strongest correlation is also with the Methylobacteriaceae (r^2^=0.341; **Supplemental Table 3**), reinforcing the idea that this clade is one of the few with a robust effect from maize genotype.

Unfortunately, we could identify no significant GWAS hits for Methylobacteria at any clade level with an empirical p-value cutoff of p≤0.01. The first PC of Weighted UniFrac analysis did show a hit on chromosome 7 (7:165,427,159, maize genome AGPv3 coordinates) as part of a cluster of metabolic GWAS hits; see the next section for more details on this cluster. If, however, we relax the p-value threshold to empirical p≤0.05, we do identify hits for two separate Methylobacteria OTUs: OTU 575956 (unknown species) at 8:8,617,635 with an empirical p-value of 0.04, and OTU 591699 (*Methylobacterium adhaesivum*) at 6:31,420,753 with an empirical p-value of 0.05. The hit on chromosome 8 is in GRMZM2G396541, which is annotated as leucine-rich repeat extensin-like protein 5, a protein that in Arabidopsis is implicated in cell wall formation (UniProt:Q9SN46). GRMZM2G396541 is also annotated as sulfated surface glycoprotein 185, which is involved in the extracellular matrix of *Volvox* (Hallmann 2003). Meanwhile, the hit on chromosome 6 is in an ABC transporter of unknown function (GRMZM2G099619). Although these annotations could logically have a role in affecting the leaf surface microbiome, the high empirical p-value for these hits means they should be interpreted with caution.

### Host genetics may influence phyllosphere metabolism more than taxonomy

It is increasingly thought that the metabolic functions of a microbial community are more important than its taxonomic identity (Coles et al. 2017; Louca et al. 2016). That is, communities which appear very different based on bacterial OTUs could actually appear quite similar when measured by the metabolic functions present. Along these lines, our analyses showed higher and more significant heritability for inferred metabolic functions than for diversity metrics or individual OTUs. Twenty metagenome terms had empirical p-values <0.0001, meaning zero out of 10,000 random permutations showed a heritability that high or higher (**Supplemental Table 7**). Unfortunately, many of these terms are poorly characterized. Of the ones with assigned functions, some of the most heritable include alkyldihydroxyacetonephosphate synthase (K00803, involved in ether lipid metabolism; h^2^=0.70), urea carboxylase (K01941, h^2^=0.70), nitrotoluene degradation (h^2^=0.68), a tRNA amidotransferase (K02433; h^2^=0.67), a circadian clock protein (K08481; h^2^=0.66), NifD nitrogenase (COG2710, involved in nitrogen fixation; h^2^=0.64), a glycan biosynthesis gene (K05946; h^2^=0.61), ureidoglycine aminohydrolase (K14977, involved in purine catabolism; h^2^=0.59), and isoquinoline 1-oxidoreductase (K07302 and K07303 for the alpha and beta subunits, respectively, and involved in isoquinoline degradation; h^2^=0.53 and 0.52). Some of these functions make sense in terms of phyllosphere biology, such as a protein likely involved in glyoxylate utilization (COG3257; h^2^=0.60), and two proteins implicated in DNA repair (COG1059 and COG1743; h^2^=0.50 and 0.49, respectively).

A Fisher’s Exact test for enrichment of terms among the highly heritable (p≤0.001) terms identified four higher-level terms that are significantly enriched (**Supplemental Table 8**): “Methane metabolism”, “Nitrotoluene degradation”, “Glyoxylate and dicarboxylate metabolism”, and “Secretion system.” Both methane metabolism and glycoxylate metabolism involve the metabolism of short (1- or 2-carbon) carbon compounds. These compounds are known to be an important energy source for phyllosphere bacteria, especially Methylobacteria (Steven E. Lindow and Brandl 2003). Nitrotoluenes are not normally considered part of the plant environment but are involved in the manufacture of agricultural chemicals (Dugal 2005), so the enrichment of this category may reflect management practices. Interestingly, nitrotoluene degradation was the only higher-level term that was both statistically enriched (p=0.01 after FDR correction) and significantly heritable on its own (h^2^=0.677, p<0.0001), making it among the most robust hits in the dataset. Finally, membrane secretion systems could be involved in many phyllosphere-important processes, including intercellular signaling, manipulation of the host, or biofilm production, but this dataset lacks the detail to know which one(s) may be most significant.

The fact that only four higher-level metabolic terms were found significant by Fisher’s Exact test and none by Annotation Enrichment Analysis (which was designed specifically to correct for biases in Fisher’s Exact test applied to hierarchical metabolism enrichment) indicates that these results should be taken with caution. This is especially true since these results are based off PICRUST-imputed metagenome content instead of actual metagenome data. For now, these results are suggestive of what processes might be significantly controlled in the phyllosphere, but further work will be needed to confirm them.

### Few phyllosphere traits show robust genome-wide associations

Our inclusion of flowering time as a positive control confirms that our analysis pipeline is capable of identifying robust, biologically relevant hits. However, the small number of hits for phyllosphere traits (most with 0 or 1 at empirical p≤0.01) indicates that these traits are more difficult to map than flowering time. Several factors could explain this. Even the most heritable phyllosphere traits have h^2^ values around 0.7, compared to 0.92 for flowering time, so that the phyllosphere traits have intrinsically lower power for mapping. They may also have a more distributed genetic architecture, or at least fewer large-effect alleles, so that each individual locus explains less variance and is harder to identify. This may explain why nitrotoluene degradation, one of the most robustly associated metagenome traits, has no GWAS hits.

Of the hits we do identify, many are close to or within known genes. Unfortunately, the majority of these genes have no known function, making interpretation of their association difficult. In addition to individual hits, however, we also identified two strong hit clusters on chromosomes 7 and 10 that are associated with multiple traits. The first is located at position 7:165,427,159 (maize genome AGPv3 coordinates) and includes 21 traits with a correlation-corrected score of 2.26, roughly equivalent to just over two completely independent traits mapping to the same location. The other is at 10:92,038,994 and includes 6 traits with a corrected score of 3.2.

Linkage disequilibrium (LD) analysis indicates that the hit cluster on chromosone 10 may be part of an extended haplotype, as there is at least one SNP ˜100kb away in very high LD (**Supplemental Figure 10**). This hit is close (˜65kb) to only a single gene of unknown function, GRMZM2G158141. In addition, the minor allele frequency at this location is very low (only 5 out of 102 individuals with non-missing genotypes), which may increase the odds of this being a spurious association due to stochastic sampling.

In contrast, the hit cluster on chromosome 7 is much more robust, with 25 minor alleles out of 230 individuals. It is also very high resolution, with no other SNPs in significant linkage disequilibrium within 10 megabases (**Supplemental Figure 10**). This hit is located just upstream (˜1kb) of GRMZM2G031545, a housekeeping gene (elongation factor 1-beta) involved in protein synthesis. The Maize Gene Expression Atlas (Stelpflug et al. 2016) (accessed via MaizeGDB.org) indicates that this gene is indeed a housekeeping gene with almost universally high expression across tissues; it is unknown why or how such a gene would affect the phyllosphere. We examined the relationship between GRMZM2G031545 and several of the traits associated with it using maize expression data from the same samples (Kremling et al. 2018), but the correlations were extremely weak (**Supplemental Figure 11**). Most of the other nearby genes have no known function, making it difficult to speculate what biological processes may be involved in this association.

### Conclusions and future directions

These results indicate that only a subset of bacteria in the maize phyllosphere are significantly impacted by the genetics of their host. This is in line with a prior analysis of the maize rhizosphere (Peiffer et al. 2013), where plant genotype had small but statistically significant effect on the rhizosphere community. In our analysis, we find robust signals for the host plant influencing a few key taxa and traits, including the abundance of Methylobacteria and the metabolism of short-chain carbon molecules and nitrotoluene.

Based on the results of the GWAS analysis, this panel is likely underpowered for phyllosphere traits. Although a panel of this size (n=246 individuals after filtering) is adequate for traits with several strong QTL, like flowering time (**Figure 7**), it appears that larger panels will be needed to map most phyllosphere-related traits. The lower heritabilities of these traits are partly responsible for this lack of power. Phyllosphere traits may also be controlled by many genes of small effect, making it more difficult to identify any individual locus as having an effect on the phenotype. We would therefore recommend including at least 350 diverse lines in future panels, and up to 400 or 500 lines if resources permit.

Although we tried to be thorough in our analysis, no single experiment can fully describe the entire phyllosphere system. There are several lines of inquiry that could complement this analysis, including looking at different phyllosphere communities (such as comparing abaxial versus adaxial surfaces) looking at different leaves down the plant or at different developmental stages, examining stalk-surface communities, etc. Above-ground endophytic communities may also have a closer relationship with host genetics than the epiphyte communities we focused on. (Although our sampling protocol did not exclude endophytes, if present they appear to be a minority of the dataset.) One could also explore if host genetics can modify the evolution of phyllosphere communities over time, since these communities are known to change over the season (Manching et al. 2018). Finally, it will be important to expand these analyses to non-bacterial members of the phyllosphere (fungi, oomycetes, etc.) to further understand phyllosphere community assembly and function.

An important take-home from our analysis is that the phyllosphere community is likely amenable to manipulationg through farmer management. This would make phyllosphere manipulation much faster and easier to accomplish than if new plant varieties had to be bred with specific genetic characteristics. A full systems-level understanding of the phyllosphere community will be necessary to understand what management practices may be beneficial to the community, and how the community as a whole could be harnessed to improve global production.

## Acknowledgements

We would like to thank members of the Buckler lab for help collecting samples (especially Shu-Yun Chen, Mei-Ksui Su, Jeremy Pardo, Nicholas Lepak, and Joshua Budka), the Genomic Diversity Facility at Cornell for help with automation, and members of both the Buckler and Wallace labs for commentary and discussions during preparation. We also thank William A. Walters for help in designing the 16s rRNA amplification. This work was supported by the University of Georgia, the USDA-ARS, and NSF grant ISO-1238014.

## Supplemental Data

- **Supplemental Figures 1-11**
- **Supplemental File 1: Bacterial diversity of the maize phyllosphere.** This file contains the original Krona plot used to generate **Figure 1**. Since these plots are interactive, it is provided to allow others to more easily explore the abundance of different taxa in the phyllosphere.
- **Supplemental Table 1: Statistical significance of alpha-diversity cofactors**. The seven non-redundant alpha diversity metrics (see Methods) were tested for correlations with different technical confounders that could drive the data. Raw p-values are shown, with the one association that is still significant after Bonferroni correction being bolded.
- **Supplemental Table 2: Statistical significance of beta-diversity cofactors.** A MANOVA analysis was conducted to identify technical cofactors that could confound sample beta diversity. The resulting p-values for each test are shown, with the most significant one per metric bolded.
- **Supplemental Table 3: Correlations between beta diversity principal coordinates and bacterial taxa.** Correlations were computed between the first 5 PCs of each beta diversity metric and all bacterial clades in our analysis; the top 10 clades are shown for each PC.
- **Supplemental Table 4: Alpha diversity heritability.** Narrow-sense heritability estimates for all tested alpha diversity measures and their resulting empirical p-value. No terms are significant.
- **Supplemental Table 5: Beta diversity heritability.** Narrow-sense heritability estimates for all tested beta diversity measures and their resulting empirical p-value. Significant terms are bolded.
- **Supplemental Table 6: OTU heritability.** Narrow-sense heritability estimates for all tested OTUs and higher-level taxonomic clades are shown with their resulting empirical p-value. Significant clades are bolded.
- **Supplemental Table 7: Predicted metagenome heritability.** Narrow-sense heritability estimates for all predicted metagenome functions are shown with their resulting empirical p-value. Significant terms are bolded, and high-heritabiltiy terms that are probably artifacts are indicated as such (see **Supplemental Figure 8**).
- **Supplemental Table 8: Enrichment of heritable metagenome functions.** The statistical enrichment of metagenome terms is shown along with the p-value from Fisher’s Exact Test (raw, Bonferroni-corrected, and FDR-corrected) and from Annotation Enrichment Analysis (REF). Terms that are significant after FDR correction are bolded.
- **Supplemental Table 9: Genome-wide association hits.** All hits from genome-wide association analysis are shown for all tested traits. (Traits not in this table had no significant hits.) The table includes the trait, marker name and position, raw and empirical p-values, clustering results, and nearest gene for all hits with empirical p-values ≤ 0.10, although only those with p ≤ 0.01 are actually considered significant.

Supplemental tables 3-9 are provided as a single zipped archive.

